# SARS-CoV-2 spike variants differ in their allosteric response to linoleic acid

**DOI:** 10.1101/2022.04.21.489022

**Authors:** A. Sofia F. Oliveira, Deborah K. Shoemark, Andrew D. Davidson, Imre Berger, Christiane Schaffitzel, Adrian J. Mulholland

**Affiliations:** Centre for Computational Chemistry, School of Chemistry, University of Bristol, Bristol BS8 1TS, UK; School of Chemistry, University of Bristol, Bristol BS8 1TS, UK; School of Biochemistry, University of Bristol, Bristol BS8 1TD, UK; School of Cellular and Molecular Medicine, University of Bristol, University Walk, Bristol, BS8 1TD, UK; Max Planck Bristol Centre for Minimal Biology, School of Chemistry, Bristol BS8 1TS, UK

## Abstract

The SARS-CoV-2 spike protein contains a fatty acid binding site, also found in some other coronaviruses (e.g. SARS-CoV), which binds linoleic acid and is functionally important. When occupied by linoleic acid, it reduces infectivity, by ‘locking’ the spike in a less infectious conformation. Here, we use dynamical-nonequilibrium molecular dynamics (D-NEMD) simulations to compare the response of spike variants to linoleic acid removal. These simulations show that the fatty acid site is coupled to functional regions of the protein, some of them far from the site (e.g. in the receptor-binding motif, N-terminal domain, the furin cleavage site located in position 679-685 and the fusion peptide-surrounding regions) and identify the allosteric networks involved in these connections. Comparison of the response of the original (‘Wuhan’) spike with four variants: Alpha, Delta, Delta plus and Omicron BA.1 show that the variants differ significantly in their response to linoleic acid removal. The allosteric connections to the fatty acid site on Alpha are generally similar to the original protein, except for the receptor-binding motif and S71-R78 region which show a weaker link to the FA site. In contrast, Omicron is the most affected variant exhibiting significant differences in the receptor-binding motif, N-terminal domain, V622-L629 and the furin cleavage site. These differences in allosteric modulation may be of functional relevance, e.g. in differences in transmissibility and virulence. Experimental comparison of the effects of linoleic acid on different variants is warranted.

## Introduction

The spike glycoprotein, which is located on the surface of the virion, mediates SARS-CoV-2 entry into host cells by binding primarily to the receptor angiotensin-converting enzyme 2 (ACE2) (*1, 2*) as well as to other receptors, such as neuropilin-1 (*3, 4*) and potentially nicotinic acetylcholine receptors (*5, 6*). The spike ectodomain (Figure 1A and Figure S2) contains the domain that directly binds to the human receptors (named receptor-binding domain, RBD) as well as all the machinery needed to fuse the host and viral membranes, including the fusion peptide (FP) (*7–9*). The SARS-CoV-2 spike contains two proteolytic cleavage sites (*7*): a furin cleavage site located at the S1/S2 junction (residues N679-R685), which distinguishes SARS-CoV-2 from other betacoronavirus spike proteins and affects the overall stability of the protein and modulates infectivity (*10–12*); and, a cleavage site in the S2 subunit (Figure S2) that releases the fusion peptide (*10*). A free fatty acid (FA) binding site was also discovered in the SARS-CoV-2 spike by members of this team in late 2020 (*13*). Subsequently, equivalent FA sites have been identified in other closely related spikes (e.g. (*13–15*)), including in the pangolin coronavirus (*14*) and the original SARS-CoV (*13*). This discovery opened the door for potential new spike-based therapies based on free fatty acids. The FA pocket is located at the interface between two neighbouring RBDs on adjacent monomers in the homotrimeric spike (*13*) (Figure 1A). This hydrophobic pocket is formed by two RBDs, with one RBD providing the aromatic and hydrophobic residues to accommodate the FA hydrocarbon tail and the RBD of the adjacent chain providing the polar (Q409) and positively charged (R408 and K417) residues that bind the FA carboxylate headgroup (*13*) (Figure 1B). The presence of linoleic acid (LA) in the FA pocket stabilises the locked spike conformation (in which the receptor-binding motifs are buried inside the RBD trimer, rendering them inaccessible for binding to ACE2), reducing infectivity (*13*).

**Figure 1.**
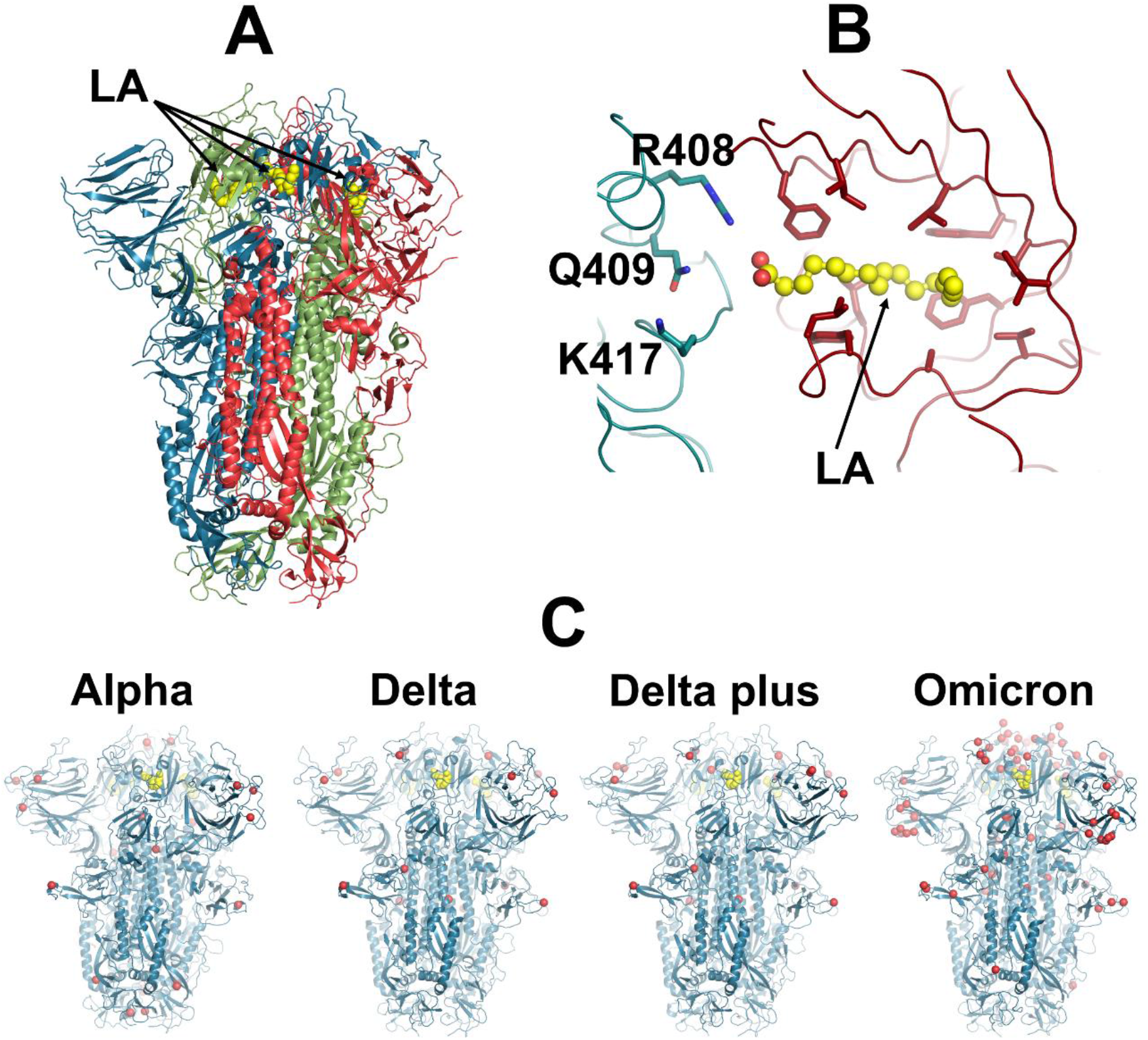
Ectodomain of the SARS-CoV-2 spike trimer with linoleic acid (LA) bound to the fatty acid-(FA) binding site. (**A**) Three-dimensional structure of the complex of the locked conformation (in which all receptor-binding motifs (RBMs) are occluded) of the ectodomain of the SARS-CoV-2 spike trimer, with LA bound (PDB code: 6ZB5) (*13*). The spike protein is a homotrimer (*7*): each monomer is shown in a different colour: green, red and blue. LA molecules are highlighted with yellow spheres. Note that each FA binding site is located at the interface between two neighbouring monomers and is formed by residues from two adjacent receptor-binding domains. (**B**) Detailed view of the FA binding site: this pocket is lined by hydrophobic and aromatic residues, and the LA acidic headgroup is located near R408, Q409 and K417. (**C**) Models used as starting points for the equilibrium MD simulations of the Alpha, Delta, Delta Plus and Omicron variants (*16*). The yellow spheres represent the LA molecules. The red spheres highlight the position of mutations, deletions and insertions in the four simulated variants.

Biomolecular simulations have provided molecular level insight into SARS-CoV-2 spike structure and dynamics, uncovering the effect of mutations, predicting interactions and revealing allosteric connections in the protein (e.g. (*3, 13, 17–30*)), including providing insight into the complex role of the FA site (*13, 17, 29, 30*). Equilibrium molecular dynamics (MD) simulations indicated persistent and stable interactions between LA and the spike trimer (*13, 17*), and that binding of LA rigidifies the FA binding site (*17*). Recently, in our previous work, using dynamical-nonequilibrium molecular dynamics (D-NEMD) simulations, we showed that the FA site is allosterically coupled to functional motifs for membrane fusion or with antigenic epitopes (*29, 30*). These simulations revealed that the removal of LA from the FA sites induces long-range structural responses in the receptor-binding motif (RBM), N-terminal domain (NTD), furin cleavage site and FP-surrounding regions (*29, 30*). D-NEMD simulations have also highlighted different allosteric and dynamical behaviours between the original (also known as ‘Wuhan’ or early 2020) spike and the D614G and BriSΔ (a variant containing an eight amino-acid deletion encompassing the furin recognition motif and S1/S2 cleavage site) variants (*29, 30*).

Here, we use D-NEMD simulations (*31–34*) to analyse the response of four spike variants to LA removal. We compare the structural and dynamical responses of the Alpha (B.1.1.7), Delta (B.1.617.2), Delta Plus (B1.617.2-AY1) and Omicron BA.1 (B.1.1.529) variants with the original (‘Wuhan’, early 2020) spike. Alpha was first detected in the UK in late 2020 and was largely responsible for the surge in cases in the winter of 2020/21 (*35–37*). This variant contains seven spike mutations and three deletions (notably L18F, H69Δ, V70Δ, Y144Δ, N501Y, A570D, P681H, T716I, S982A, D1118H) and is more transmissible than the original virus (*35–37*). Delta was initially identified in India in late 2020 and was, until recently, the dominant strain globally (https://cov-lineages.org/global_report_B.1.617.2.html). It harbors six mutations and three deletions in the spike (T19R, E156Δ, F157Δ, R158Δ, L452R, T478K, D614G, P681R and D950N) and shows enhanced transmissibility compared to the previous variants (*38*). By the middle of 2021, a variant of Delta with the K417N mutation (nicknamed as ‘Delta Plus’) was identified in Nepal and quickly spread to the rest of the world (https://cov-lineages.org/lineage.html?lineage=AY.1). Note that although Delta Plus spread worldwide, it did not displace Delta in the same way the Omicron did (for more details, see SARS-CoV-2 sequences by variant, Apr 14, 2022 (ourworldindata.org)). The Omicron BA.1 variant (hereafter referred to as Omicron) was initially detected in South Africa in November 2021 (*39, 40*) and spread rapidly globally and has become the dominant variant in many countries (see SARS-CoV-2 sequences by variant, Apr 14, 2022 (ourworldindata.org)). This variant includes up to 40 mutations, deletions and insertions on the spike (A67V, H69Δ-V70Δ, T95I, G142D, V143Δ, Y144Δ, Y145Δ, N211Δ, L212I, D215E, PED insertion in position 210-212, G339D, S371L, S373P, S375F, K417N, N440K, G446S, S477N, T478K, E484A, Q493R, G496S, Q498R, N501Y, Y505H, T547K, D614G, H655Y, N679K, P681H, N764K, D796Y, N856K, Q954H, N969K and L981F) (*41*). Omicron is more transmissible than Delta and any other preceding variants but apparently causes less severe disease than previous strains (e.g. (*42–44*)).

D-NEMD simulations (*34*) are emerging as a practical technique for identifying structural communication pathways in various biomolecular systems (*45–48*). For the original (‘Wuhan’) spike, this approach identified allosteric connections between the fatty acid site and functionally important regions of the protein (including the RBM, furin cleavage site and FP-surrounding regions) and mapped the networks that connect them to the FA site (*29, 30*). D-NEMD simulations have also been used to investigate the effects of pH on the Delta spike (*26*).

Hundreds of D-NEMD simulations were performed to analyse the response of the original spike and of the Alpha, Delta, Delta Plus and Omicron variants to LA removal. The models used here for the original, Alpha, Delta, Delta plus and Omicron were taken from ref. (*16*) and were built using the cryo-EM structures of the locked ectodomain of the spike with LA bound (*13*) and the NTD of the NOVAVAX structure (*15*) (as it had better-defined loops for the NTD than previous structures (*13*)). We compare the structural and dynamical responses of the variants to the original protein. These simulations investigate how mutations, insertions and deletions affect the allosteric pathways connecting the FA site to the rest of the protein. In the D-NEMD approach, multiple configurations extracted from equilibrium simulations are used as starting points for nonequilibrium simulations, through which the effect of a perturbation can be studied (*34*). The time-dependent response of the protein to the perturbation is extracted by directly comparing equilibrium and nonequilibrium trajectories at equivalent points in time (Figure S1). Equilibrium trajectories for the locked form of the non-glycosylated and uncleaved (no cleavage at the furin recognition site) ectodomain of the original, Alpha, Delta, Delta Plus and Omicron spike bound with LA were taken from ref. (*16*). For each system, 90 short (5 ns) D-NEMD simulations were carried out starting from conformations extracted from the equilibrium simulations (Figure S1). In this work (and similarly to our previous work (*29, 30*)), the perturbation introduced to the protein was the instantaneous deletion of all LA molecules from the FA sites. Glycans are crucial to the biological functions of the spike, being important for shielding (*18*) and infection (e.g. by altering the dynamics of RBD opening (*18, 27*)). To investigate e.g. interactions with receptors, including the glycan shield is important (e.g. (*6, 18*)). The aim here was to compare the response of the variant spike proteins. The allosteric communication networks and the response of the protein to an internal structural perturbation (here, LA removal) are unlikely to be qualitatively altered by the presence of glycans, as they cover the outside of the protein. The cryo-EM structure of the original spike in a locked conformation with LA bound (*13*) only contains glycans on the exterior.

In the D-NEMD simulations, the instantaneous deletion of the LA molecules from the FA sites prompts the structural response of the protein as it adapts to LA removal. Note that the D-NEMD simulations are not intended to model the physical process of LA binding or dissociation. The perturbation used here, which was the same as in our previous work (*29, 30*), is designed to induce a rapid response and force signal transmission within the protein, hence allowing mapping of the mechanical and dynamical coupling between the structural elements involved in this process. The time evolution of the response is extracted using the Kubo-Onsager relation (*31–34*). Multiple D-NEMD simulations are performed and compared with the equilibrium trajectories to identify the protein’s structural response. This response is averaged over multiple trajectories, removing noise (*31–34*). Given that the same perturbation (LA removal) is used for all systems, the structural response of the different variants can be directly and meaningfully compared. Note that the D-NEMD approach allows for the statistical significance of the structural rearrangements to be determined (Figures S4-S8) (*34*). We focus on the differences shown here to be significant.

The D-NEMD simulations reveal the complex cascade of conformational changes induced by LA removal and identify the pathways through which these changes propagate within the protein. The structural response quickly propagates from the FA site to specific and well-defined regions of the spike, in all the variants (Figure 2 and S9). Several functional motifs, including the regions surrounding the FP, show significant structural responses in all variants (Figure 2). In the systems simulated, LA removal induces a conformational response in the FA pocket, which contracts due to movement of the hydrophobic and aromatic sidechains that line it (Figure S10). Structural changes are then rapidly transmitted to the RBD and NTD, and then to V622-L629, the furin cleavage site, and residues surrounding the FP (Figure S9). As can be seen in Figures 2, 3 and S9, 0.1 ns after LA removal, significant structural rearrangements can already be observed in the RBD, mainly in the RBM, and in specific regions of the NTD, such as S71-R78, H146-E156 and L249-G257. The RBM contains the residues that directly interact with the host ACE2 receptor (*8, 49*) and is a known target for neutralising antibodies (*50–52*). The NTD is also a major target for neutralising antibodies (e.g. (*53–56*)). In particular, the H146-E156 and L249-G257 segments were shown to directly mediate the interaction between spike and specific antibodies (e.g. 4A8 monoclonal antibody) (*54*). The S71-R78 region is an antigenic epitope and has also been suggested to be involved in binding to other receptors besides ACE2 (*57*).

**Figure 2.**
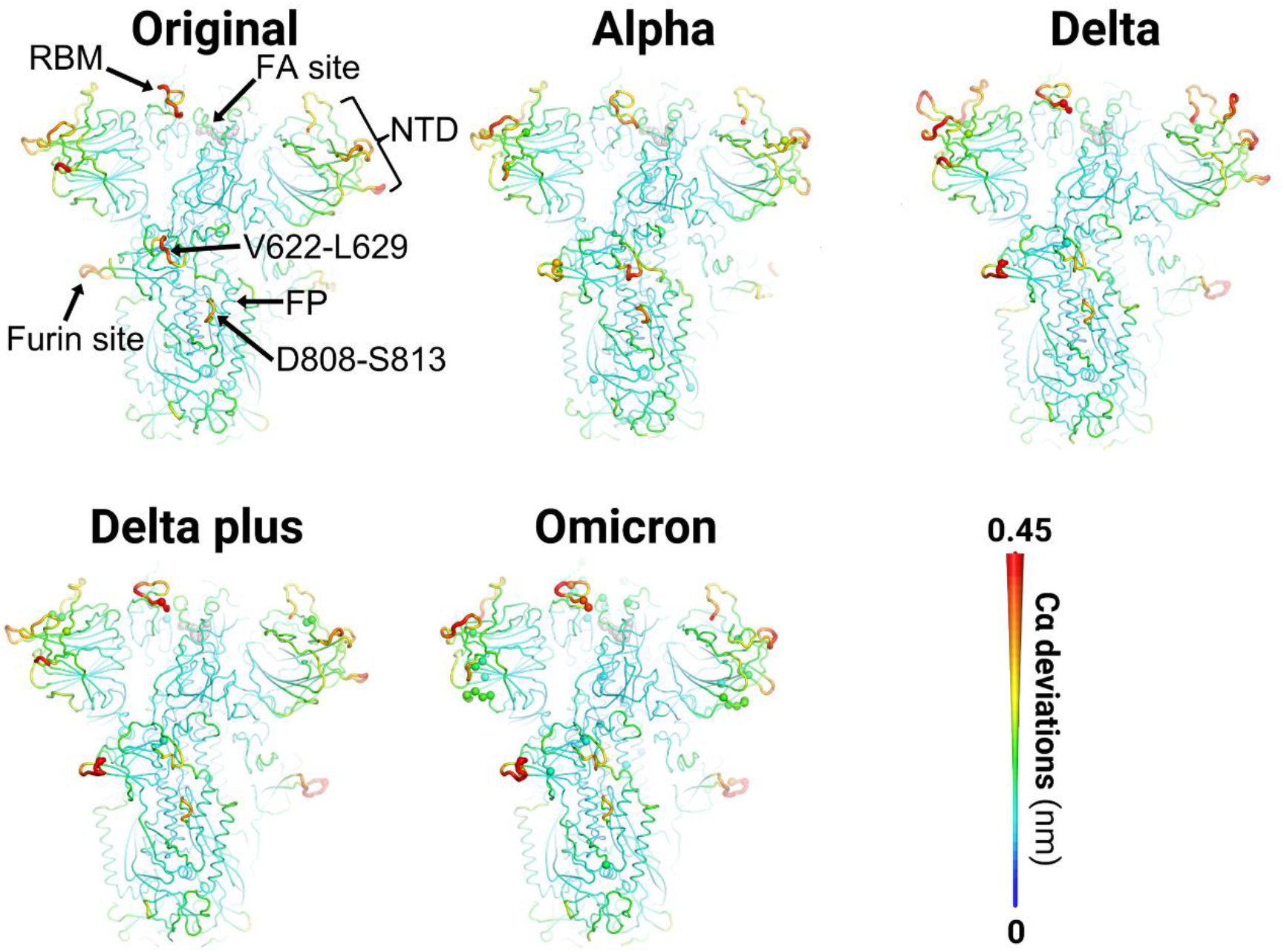
Structural response of the original, Alpha, Delta, Delta plus and Omicron variants. Mapping of the average Cα-positional deviations after LA removal from the FA binding sites. The average Cα deviations between the D-NEMD apo and equilibrium LA-bound simulations were calculated for each residue. The final deviation values correspond to the average obtained over the three chains of the trimer and over the 90 pairs of simulations (Figure S3). The Cα average deviations after 5 ns are mapped onto the starting structures for equilibrium simulations of each variant. Structure colours and cartoon thickness both indicate the average Cα-positional deviation values (indicated in the scale on the right). The grey spheres highlight the FA binding site. The other spheres pinpoint the position of mutations.

**Figure 3.**
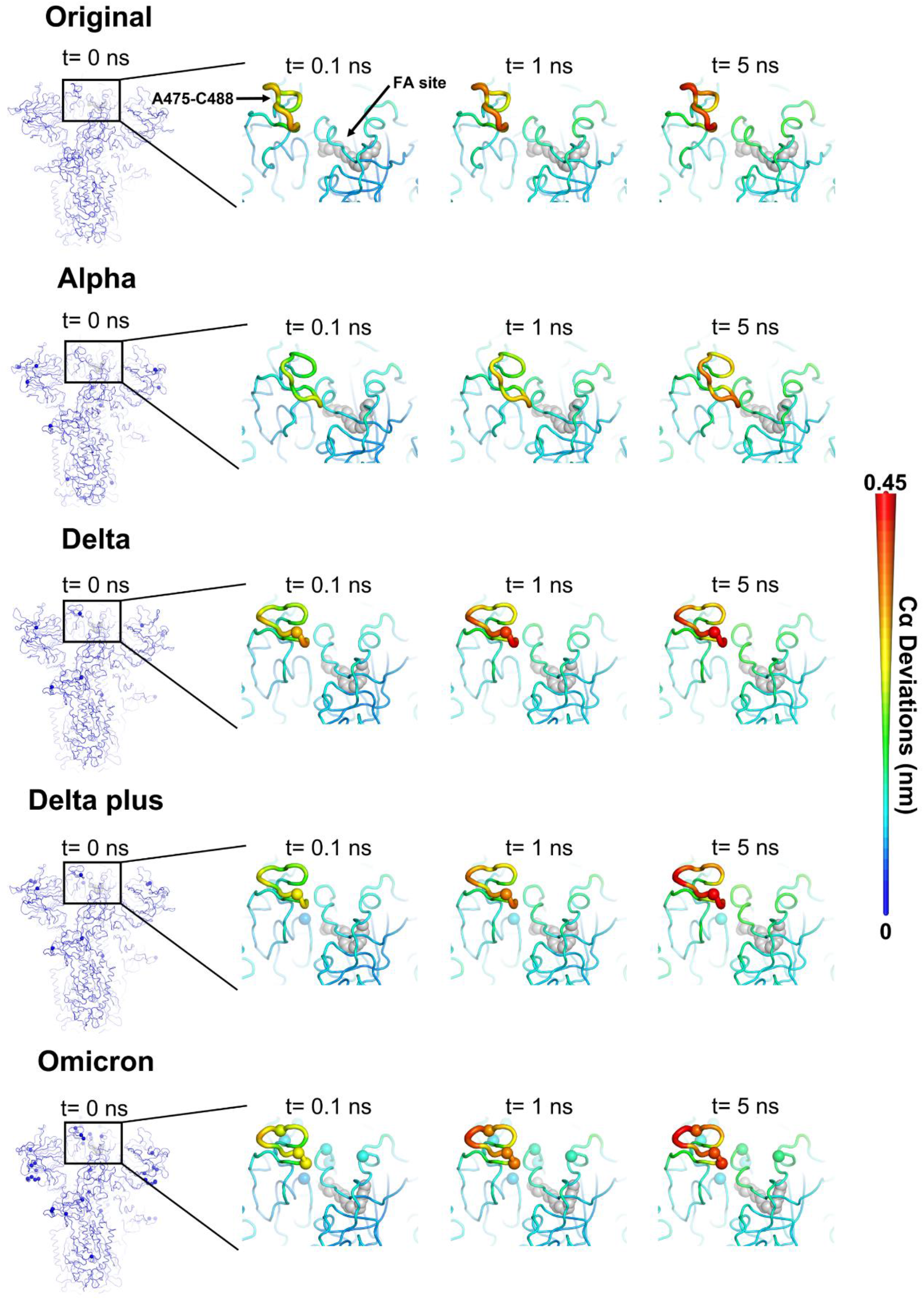
Structural responses of the RBD in the original, Alpha, Delta, Delta plus and Omicron variants. The Cα average deviations are mapped onto the starting structures for equilibrium simulations of each variant. Structure colours and cartoon thickness both indicate the average Cα-positional deviation values (indicated in the scale on the right). The grey spheres highlight the FA binding site. The other spheres show the position of mutations. For more details, see the legend in Figure 2.

The networks that connect the FA site to functional regions of the spike are similar in all the variants, thus apparently are conserved (Figure S9). Similarly to what was observed for the original spike in our previous work (*29*), the S366-A372 and R454-K458 segments transmit the structural changes from the FA site to the RBM (Figure S11), while the P337-A348, W353-I358 and C166-P174 regions mediate signal propagation to the NTD (Figure S12) in all variants. Signal transmission to the S1/S2 interface and S2 subunit occurs via the C525-K537, F318-I326, and L629-Q644 regions (Figure S13).

The different variants exhibit similar allosteric networks, but there are significant differences in their responses (Figures 3–5). Comparing the time evolution of the protein’s response shows significant differences in behaviour between the variants (Figure S9). The variants differ from the original spike in the amplitude of the structural response to the LA removal and the rate at which these rearrangements propagate within the protein structure (Figures 3–5). Functionally important regions are affected to significantly greater or lesser extents in the variants, as detailed below.

**Figure 4.**
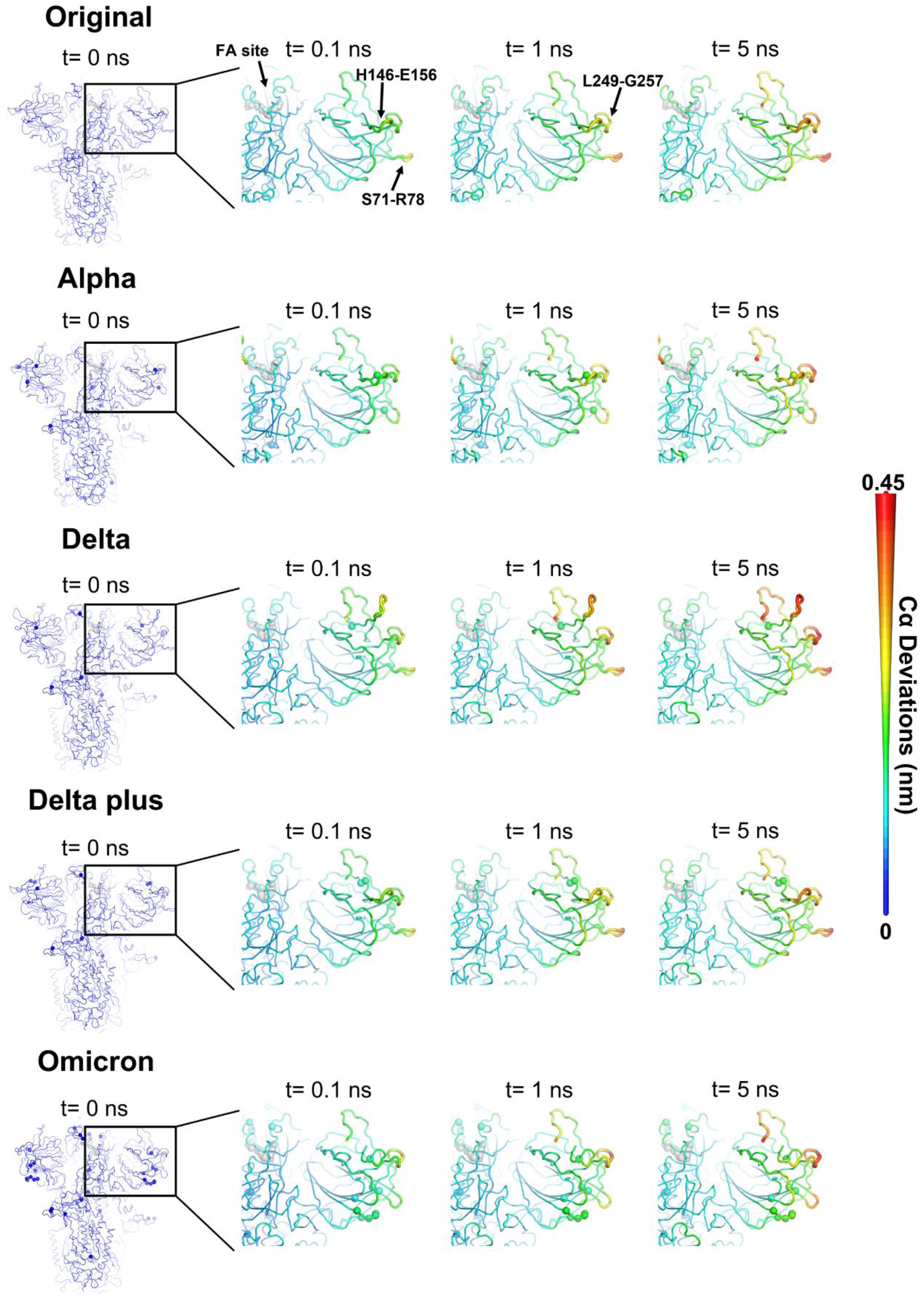
Structural response of the NTD in the original, Alpha, Delta, Delta plus and Omicron spike variants. The Cα average deviations are mapped onto the starting structures for equilibrium simulations of each variant. Structure colours and cartoon thickness both indicate the average Cα-positional deviation values (indicated in the scale on the right). The grey spheres highlight the FA binding site. The remaining spheres pinpoint the position of mutations. For more details, see the Figure 2 legend.

**Figure 5.**
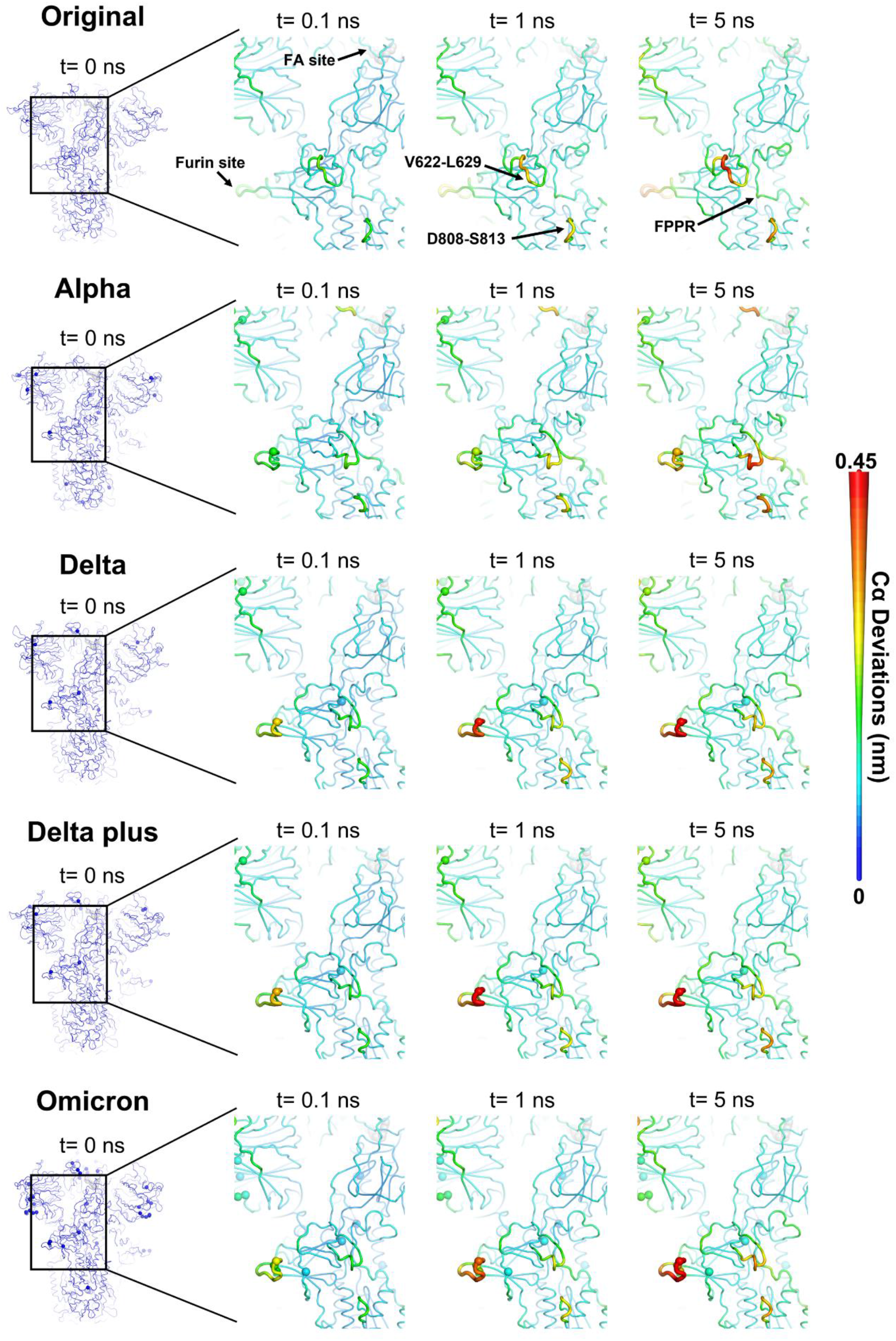
Structural response of the furin cleavage site and FP-surrounding regions in the original, Alpha, Delta, Delta plus and Omicron variants. The Cα average deviations are mapped onto the starting structures for equilibrium simulations of each variant. Both structure colours and cartoon thickness relates to the average Cα-positional deviation values (indicated in the scale on the right). The grey spheres highlight the FA binding site. The remaining spheres pinpoint the position of mutations. For more details, see the Figure 2 legend.

## Receptor binding motif

Significant variations in the response of the RBM (mainly of the A475-C488 segment) were observed between the five virus spike proteins (Figures 3 and S3). In the Alpha spike, the response of A475-C488 is notably weaker and more diffuse (the amplitude of the structural changes is lower) than the original protein (Figures 3 and S3). In contrast, Delta, Delta plus and Omicron show a larger response for the A475-C488 segment than the original spike (Figures 3 and S3). Delta, Delta plus and Omicron all have mutations in the region A475-C488, the RBD region with the largest responses to LA removal. Delta and Delta plus have a threonine-to-lysine substitution in position 478 (T478K). Omicron also contains the S477N and E484A mutations, as well as T478K. The D-NEMD simulations indicate that these mutations alter the dynamics of signal transmission from the FA site to the RBD. In Delta, Delta plus and Omicron, they amplify the allosteric coupling between the FA site and the RBM (Figures 3 and S3).

## N-terminal domain

The response of the NTD differs significantly between variants, particularly in the S71-R78, H146-E156 and L249-G257 segments (Figure 4). For example, Delta shows larger structural rearrangements in the region H146-E156 than the original protein, whereas Alpha and Omicron show reduced responses (Figures 4 and S3). These variants contain deletions and mutations either in, or in direct contact with, the H146-E156 region: Alpha contains a one-residue deletion in position 144 (Y144Δ); Delta has a three-residue deletion from position 156 to 158 (E156Δ, F157Δ and R158Δ); and Omicron harbors a three-residue deletion from position 143 to 145 (V143Δ, Y144Δ and Y145Δ). The Delta plus variant, which contains the E156Δ-F157Δ deletion plus an arginine-to-glycine mutation in position 158 (R158G), shows a similar response to the original protein. This suggests that R158G mitigates the effects of the E156Δ-F157Δ deletion.

All the variants show weaker structural responses of the S71-R78 segment than the original spike (Figures 4 and S3). The difference in this behaviour between the variants is probably due to mutations in the regions close or in direct contact with S71-R78: Alpha includes a two-residue deletion in positions 69 and 70 (H69Δ-V70Δ); Delta and Delta plus contain a threonine-to-arginine substitution in position 19 (which is located close to S71-R78); and, Omicron, as well as the H69Δ-V70Δ deletion, also contains an alanine-to-valine mutation in position 67 (A67V). It was recently found that mutations in the RBD (e.g. E484A) may be compensated for by stabilising mutations, e.g. in the NTD (H69Δ-V70Δ and G142D) and S2 domain, in the Omicron spike (*58*).

The L249-G257 region in the NTD also responds differently in the variants, with Alpha, Delta and Omicron showing enhanced structural rearrangements relative to the original spike (Figures 4 and S3). The structural response of the L249-G257 segment is modulated by the deletions and substitutions in NTD regions adjacent to this loop: L18F and Y144Δ in Alpha; T19R in Delta; and G142D and V143Δ-Y145Δ in Omicron.

## Furin cleavage site and fusion peptide proximal region

The rearrangements induced by LA removal are not restricted to the regions near the FA site, and they propagate as far as the V622-L629 segment, the furin cleavage site and the regions surrounding the FP (Figure 5 and S9). The furin cleavage site, which is located at the S1/S2 interface, shows significant differences in response to LA removal between variants (Figures 5 and S3). In Alpha, the furin cleavage site is less impacted by LA removal than in the original protein. In contrast, in Delta, Delta plus and Omicron, the furin cleavage site is more affected (Figure S14). Alpha, Delta, Delta plus, and Omicron contain residue substitutions close to the furin cleavage site. The proline residue in position 681 is mutated to histidine in Alpha and Omicron (P681H) and to an arginine in Delta and Delta Plus (P681R). In addition to P681H, in Omicron, asparagine 679 is also replaced by lysine (N679K). The extra positively charged residues near the cleavage site (P681R in Delta and Delta plus and N679K in Omicron) strengthen the allosteric connection to the FA pocket (Figure S14). The addition of flanking positively charged residues to the P681-R685 stretch has been suggested to improve proteolytic processing (*59*).

The allosteric coupling between the FA site and V622-L629 is substantially weaker in the variants containing the D614G mutation (Delta, Delta Plus and Omicron): the structural changes are smaller than the original spike (Figures 5 and S14). D614G significantly reduces signal propagation and allosteric coupling between the FA site and V622-L629. The D614G mutation is located at the interface between two monomers, where it disrupts the trans-interface salt-bridge and hydrogen bond networks (e.g. (*60, 61*)) and alters the dynamics of this region (e.g. (*29*)). The D614G substitution has been shown to increase transmission, infectivity and viral fitness (e.g. (*60, 62–67*)). Our results here indicate that it may have a role in limiting the allosteric effects of the FA site.

The regions surrounding the fusion peptide, notably D808-S813 and the FPPR, are also affected by LA removal (Figures 5 and S3). The response of D808-S813 is similar between all simulated variants (Figure S14). This segment is located upstream of the FP and immediately preceding the S2’ protease cleavage site. The S2’ site is essential for infection (e.g. (*10*)), and its cleavage is mediated by the transmembrane protease serine 2 (TMPRSS2) after spike binding to ACE2 (e.g. (*10, 68*)). Finally, the FPPR shows reduced response in all the variants compared to the original protein (Figure S14). This diminished response in Alpha, Delta, Delta plus and Omicron indicates a weakened allosteric connection of the FPPR to the FA site. The results here show that the mutations occurring in or close to the furin cleavage site and V622-L629, such as D614G (in Delta, Delta plus and Omicron), H655Y and N679K (in Omicron), P681H (in Alpha and Omicron) or P681R (in Delta and Delta plus), alter the allosteric networks connecting the FA site to the regions surrounding the FP, particularly the FPPR.

## Conclusions

Our findings show that SARS-CoV-2 variants differ significantly in their allosteric response to fatty acid binding (Table 1). These differences are of potential functional importance in the regulation of viral infectivity by LA. It may also have implications for efforts to target the FA site with natural, repurposed or specifically designed ligands (*17*).

**Table 1.** Summary of the structural response of functional regions of the spike in Alpha, Delta, Delta plus and Omicron compared to the original (‘Wuhan’) SARS-CoV-2 spike. The ↑ and ↓ arrows indicate an increase or decrease in the response relative to the original protein. The light blue, sky blue and dark blue colours represent a decrease of <5%, 5-10% and >10% in the amplitude of the response relative to the original protein, respectively. The yellow and red colours represent an increase of 5-10% and >10% in the amplitude of the response relative to the original protein. The white colour indicates that no significant change (n.s.c.) in the amplitude of the response relative to the original protein is observed.

|  | Variants |  |  |  |
| --- | --- | --- | --- | --- |
|  | Alpha | Delta | Delta plus | Omicron |
| **NTD** |  |  |  |  |
| S71-R78 | ↓ | ↓ | ↓ | ↓ |
| H146-E156 | ↓ | ↑ | n.s.c. | ↓ |
| L249-G257 | ↑ | ↑ | ↓ | ↑ |
| **RBM** | ↓ | ↑ | ↑ | ↑ |
| **Furin cleavage site** | ↓ | ↑ | ↑ | ↑ |
| **V622-L629** | n.s.c. | ↓ | ↓ | ↓ |
| **D808-S813** | n.s.c. | n.s.c. | n.s.c. | n.s.c. |
| **FPPR** | ↓ | ↓ | ↓ | ↓ |

The allosteric connections in Alpha are generally similar to the original (‘Wuhan’) spike protein, except for the RBM and S71-R78 (Table 1). Delta and Delta plus exhibit significant differences in the responses of the NTD, furin cleavage site and V622-L629 but not in the RBM (Table 1). Omicron, the most infectious variant simulated, displays significant changes in the response of the NTD, RBM, furin cleavage site and V622-L629 compared to the original spike protein (Table 1). In Omicron, S71-R78, H146-E156 and V622-L629 exhibit a weaker connection to the FA site, whereas the L249-G257 region, RBM and the furin cleavage site show stronger coupling to the FA site.

In Delta, Delta plus and Omicron, the allosteric connection between the FA site and the furin cleavage site is increased compared to the original spike, whereas the link to V622-L629 is diminished. Also, in all variants, FPPR displays weaker connection to the FA site (Table 1). This indicates that mutations affect how the structural rearrangements are propagated to the FPPR.

While all variants contain similar networks connecting the FA site to functional regions of the protein, there are statistically significant differences in their responses. Substitutions, insertions, and deletions affect the amplitude of the structural response of specific regions in the S1 and S2 subunits and alter the rates at which these rearrangements propagate to them. While some mutations (such as L18F, T19R, G142D, E156Δ-F157Δ, T478K or P681H/R) strengthen the links to the FA pocket, others (such as H69Δ-V70Δ, Y144Δ and D614G) abate them.

The coupling of the FA site to the NTD, in particular to the S71-R78, H146-E156 and L249-G257 segments, is greatly affected by mutations in and around those regions. While deletions around position 156 (as E156Δ-F157Δ in Delta) enhance the allosteric connection to the FA site, deletions around position 144 (as Y144Δ in Alpha and V143Δ-Y145Δ in Omicron) diminish that connection. Deletions near position 71 (as H69Δ-V70Δ in Alpha) reduce the response of S71-R78 and weaken the allosteric link to the FA site. Differences in the regions next to L249-G257 (for example, L18F in Alpha, T19R in Delta or G142D in Omicron) can amplify the structural rearrangements induced by LA removal, reinforcing the connection to the FA pocket.

The D-NEMD simulations also reveal that substitutions leading to an increase of the positive charge in the region preceding the furin cleavage site (i.e. P681R in Delta and Delta plus and N679K in Omicron) apparently reinforce the allosteric link to the FA pocket. Conversely, the D614G mutation (found in Delta, Delta plus and Omicron) significantly reduces the coupling of V622-L629 to the FA site.

Our results demonstrate that D-NEMD simulations are a valuable tool to study allostery, identify and explore the allosteric networks operating within the spike, and predict the impact of future mutations in these pathways. D-NEMD simulations provide an effective approach to characterise allosteric effects (e.g. (*29, 30*)), effects of pH changes (*26*) and other functionally important properties and will be helpful to investigate further emerging variants. We note that the simulations here compare spike proteins without glycans; as discussed above and elsewhere (*29*), the presence of glycans (covering the outside of the protein) is unlikely to qualitatively alter the internal mechanical response of the protein. The differences in allosteric response to LA between variants revealed by D-NEMD simulations here may have functional relevance and should be investigated by experiments.

## Acknowledgements

AJM and ASFO thank EPSRC (grant number EP/M022609/1), BBSRC (grant number BB/R016445/1) and ERC (Advanced Grant PREDACTED https://cordis.europa.eu/project/id/101021207) for support. MD simulations were carried out using the computational facilities of the Advanced Computing Research Centre, University of Bristol (http://www.bris.ac.uk/acrc) under an award for COVID-19 research and using the Oracle Public Cloud Infrastructure (https://cloud.oracle.com/en_US/iaas) under an award for COVID-19 research. We thank BrisSynBio, a BBSRC/EPSRC Synthetic Biology Research Centre (Grant Number:BB/L01386X/1) for funding ASFO and EPSRC via HECBIOSIM (hecbiosim.ac.uk) for providing ARCHER/ARCHER2 time through a COVID-19 rapid response call. We also thank Oracle Research for Oracle Public Cloud Infrastructure (https://cloud.oracle.com/en_US/iaas) time, the [Bristol UNCOVER Group](https://www.bristol.ac.uk/research/impact/coronavirus/research-priorities/) and the University of Bristol, for their support. C.S. and I.B. are Investigators of the Wellcome Trust (210701/Z/18/Z; 106115/Z/14/Z).

## Data availability statement

All D-NEMD simulation data (including input and trajectories files) are openly available from the MolSSI/BioExcel COVID-19 public data repository for biomolecular simulations (https://covid.molssi.org/simulations/).

## Notes

### Competing Interest Statement

The authors declare competing interests. Imre Berger and Christiane Schaffitzel report shareholding in Halo Therapeutics Ltd related to this Correspondence.

