## Supplementary Material for "SARS-CoV-2 spike variants differ in their allosteric response to linoleic acid"

#### Materials and methods

##### Equilibrium simulations

The sequence for the 'Wuhan' or early 2020 spike (hereafter named original) was taken from Uniprot P0DTC2 (https://www.uniprot.org/uniprot/P0DTC2). The sequences for Alpha (B.1.1.1.7), Delta (B.1.617.2) and Delta Plus (B1.617.2-AY1) were taken from UK-COG (https://www.cogconsortium.uk/). The sequences for Omicron BA.1 (hereafter named Omicron) were taken from https://www.gisaid.org as EPI-ISL-6640916. The Alpha variant comprises seven mutations and three deletions in each spike monomer, namely L18F, H69Δ-V70Δ, Y144Δ, N501Y, A570D, P681H, T716I, S982A, D1118H. The Delta variant contains six mutations and three deletions in the spike (T19R, E156Δ-R158Δ, L452R, T478K, D614G, P681R and D950N), whereas the Delta Plus has eight mutations and two deletions (T19R, E156Δ-F157Δ, R158G, K417N, L452R, T478K, D614G, P681R and D950N). The Omicron variant bears 40 substitutions, deletions and insertions (A67V, H69Δ-V70Δ, T95I, G142D, V143Δ-Y145Δ, N211Δ, L212I, D215E, PED insertion, G339D, S371L, S373P, S375F, K417N, N440K, G446S, S477N, T478K, E484A, Q493R, G496S, Q498R, N501Y, Y505H, T547K, D614G, H655Y, N679K, P681H, N764K, D796Y, N856K, Q954H, N969K and L981F), 15 of them located in the RBD.

The models for the non-glycosylated and uncleaved (no cleavage at the furin site located in the S1/S2 interface) locked ectodomain of the original, Alpha, Delta, Delta plus and Omicron with linoleic acid (LA) bound were taken from (*1*). The model for the original spike with LA was based on the cryo-EM structures 7JJI (*2*) and 6ZB5 (*3*). Note that both cryo-EM structures used to build the model for the original spike contains glycans on the exterior (*2*, *3*): limited glycosylation occurs in the structure 6ZB5 because it was expressed in insect cells (*3*); site-specific glycosylation analysis detected glycosylation in all 22 N-linked glycan sites in the structure 7JJI (*2*). Both cryo-EM structures (pdb codes: 6ZB5 and 7JJI) contain 2-acetamido-2-deoxy-beta-D-glucopyranose-(1-4)-2-acetamido-2-deoxy-beta-D-glucopyranose oligosaccharides and 2-acetamido-2-deoxy-beta-D-glucopyranose monosaccharides (*2*, *3*). The models for the Alpha, Delta, Delta plus and Omicron spikes with LA bound were constructed using as template the model for the original spike described above (for more details, see (*1*)).

The equilibrium simulations for the original, Alpha, Delta, Delta plus and Omicron used here as the starting point for the dynamical-nonequilibrium (D-NEMD) simulations were taken from (*1*). All MD simulations here used the same conditions and protocols as applied successfully previously (*3-6*). Each spike trimer was simulated in a box of explicit waters with 150 mM NaCl, under periodic boundary conditions as an NPT ensemble at 310 K and pH 7, as described in (*1*). Three replicate simulations, each 200 ns, were performed for each spike system using GROMACS (*7*).

#### D-NEMD simulations

D-NEMD simulations were performed to study the structural response to LA removal in the unglycosylated, uncleaved original, Alpha, Delta, Delta Plus and Omicron spike from SARS-CoV-2. Ninety short nonequilibrium simulations were carried out for each variant, in a total of 450 simulations. D-NEMD simulations have successfully been used to study allostery in various biomolecular systems (*8-12*), including the SARS-CoV-2 spike (*4, 5*). Recently, this approach was used to identify the allosteric networks connecting the fatty acid binding site to key functional motifs on the original, D614G (*4*) and BriSΔ (a variant containing an eight amino-acid deletion in the furin recognition motif and S1/S2 cleavage site) spikes (*5*).

Briefly, in the D-NEMD approach, the response of a system to an external perturbation (in this case, LA removal from the fatty acid binding sites) can be directly computed using the Kubo-Onsager relation (*13-16*) and by calculating the difference of a given property between the simulations with and without the perturbation. Subtracting the perturbed and unperturbed pairs of simulations at a given time and averaging the results over tens/hundreds of replicates allows not only the identification of the events associated with signal propagation, but also the determination of the statistical significance of the observations (*13-16*). Here and similarly to our previous spike work (*4, 5*), the perturbation was generated by the (instantaneous) removal of the LA molecules from the fatty acid (FA) binding sites. A graphical representation of the procedure used to set up the D-NEMD simulations is shown in the top panel of Figure S1.

The starting conformations for the D-NEMD simulations were obtained from the equilibrated part (from 50-200 ns) of the equilibrium LA-bound simulations of the original, Alpha, Delta, Delta Plus and Omicron systems (*1*). Conformations were taken every five ns, and in each frame, the LA molecules were (instantaneously) deleted from the FA pockets. The resulting apo system was then simulated for 5 ns (Figure S1). The simulation conditions for all the nonequilibrium simulations were identical to the equilibrium ones described in (*1*). 90 short apo D-NEMD simulations were performed for each system. For more details regarding the setup of the D-NEMD simulations, see (*4, 5*).

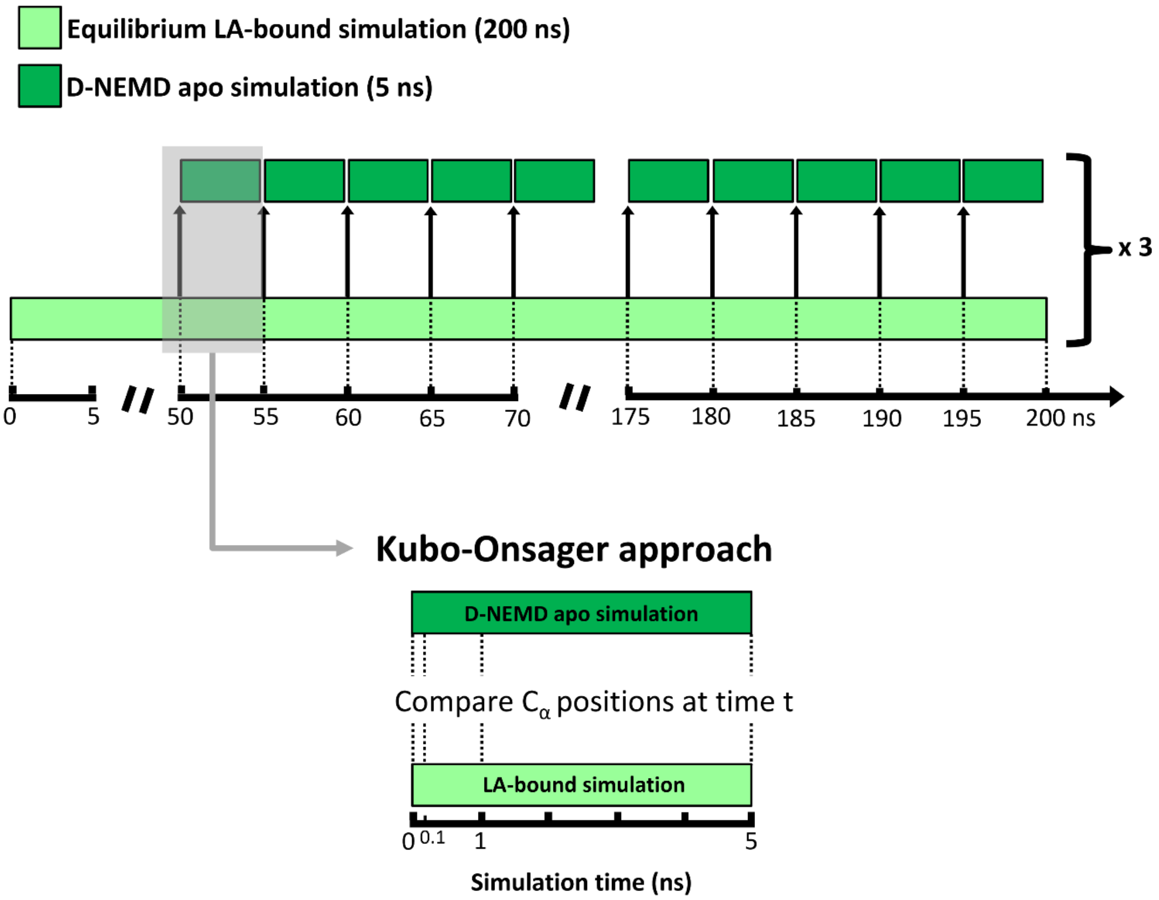


**Figure S1.** Schematic description of the procedure used to set up and analyse the D-NEMD simulations. For all systems, namely the original, Alpha, Delta, Delta Plus and Omicron spike, 3 equilibrium MD simulations, 200 ns each, were performed for the unglycosylated and uncleaved (no cleavage at the S1/S2 interface) locked spike. These equilibrium simulations (light green rectangles) were then used to generate starting structures for the short apo D-NEMD simulations (dark green rectangles). From the equilibrated part of each LA-bound simulation (from 50-200 ns), conformations were extracted every five nanoseconds, and the perturbation was introduced. Each short D-NEMD simulation was simulated for 5 ns. The Kubo-Onsager (*13-16*) approach was used to extract the response of the system to LA annihilation from the FA pockets (bottom panel). For that, for each pair of equilibrium LA-bound and D-NEMD apo simulations, the positional deviations of each Cα at equivalent times (namely 0, 0.1, 1 and 5 ns) were determined and averaged over all the simulations.

The Kubo-Onsager (*13-16*) approach was used to extract the response of the spike to LA removal (see the bottom panel in Figure S1). For each pair of unperturbed LA-bound equilibrium and perturbed apo nonequilibrium simulations, the difference in positions for each Cα was determined at equivalent points in time, namely after 0, 0.1, 1, 3 and 5 ns of simulation (Figure S3). The pairwise comparison between the positions of Cα atoms allows for the direct identification of the most important conformational rearrangements. The Cα-positional deviations between the equilibrium and nonequilibrium trajectories at each point in time were averaged over all simulations and over the three chains forming the trimer. The statistical significance of the structural changes identified here is demonstrated by the low standard error of the averages (Figures S4-S8).

### Supplementary Figures

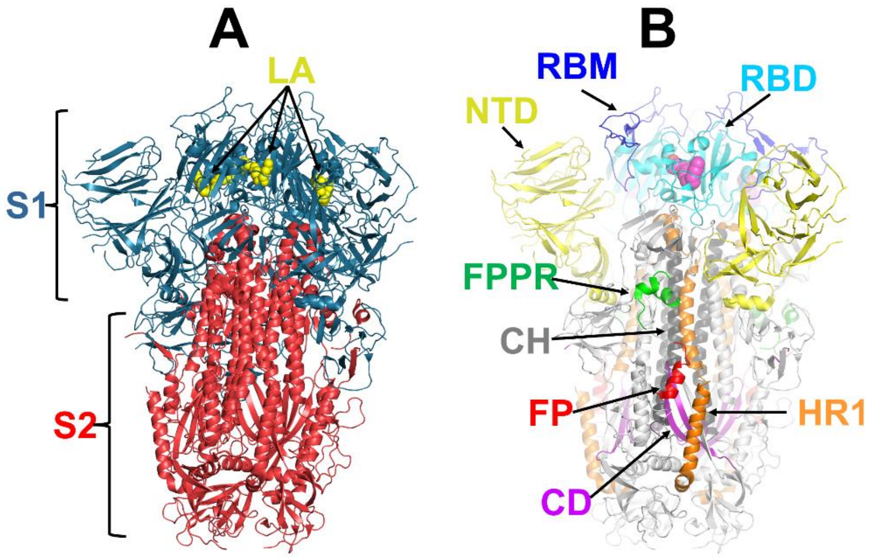


**Figure S2. (A)** Structure of the complex of the ectodomain of the SARS-CoV-2 spike trimer with linoleic acid (*3*). The S1 and S2 subunits are colored in blue and red, respectively. Linoleic acid (LA) molecules are highlighted with yellow spheres. **(B)** Structure of the ectodomain of the spike trimer (*3*) with some relevant structural motifs highlighted: N-terminal domain (NTD)-yellow; receptor-binding domain (RBD)- cyan; receptor binding motif (RBM)- blue; fusion peptide (FP)- red; fusion-peptide proximal region (FPPR)- green; heptad repeat 1 (HR1)- orange; central helix (CH)- grey; connector domain (CD)- purple. LA is shown with magenta spheres. Note that A and B represent the ectodomain of the spike trimer in the locked conformation (three closed RBDs).

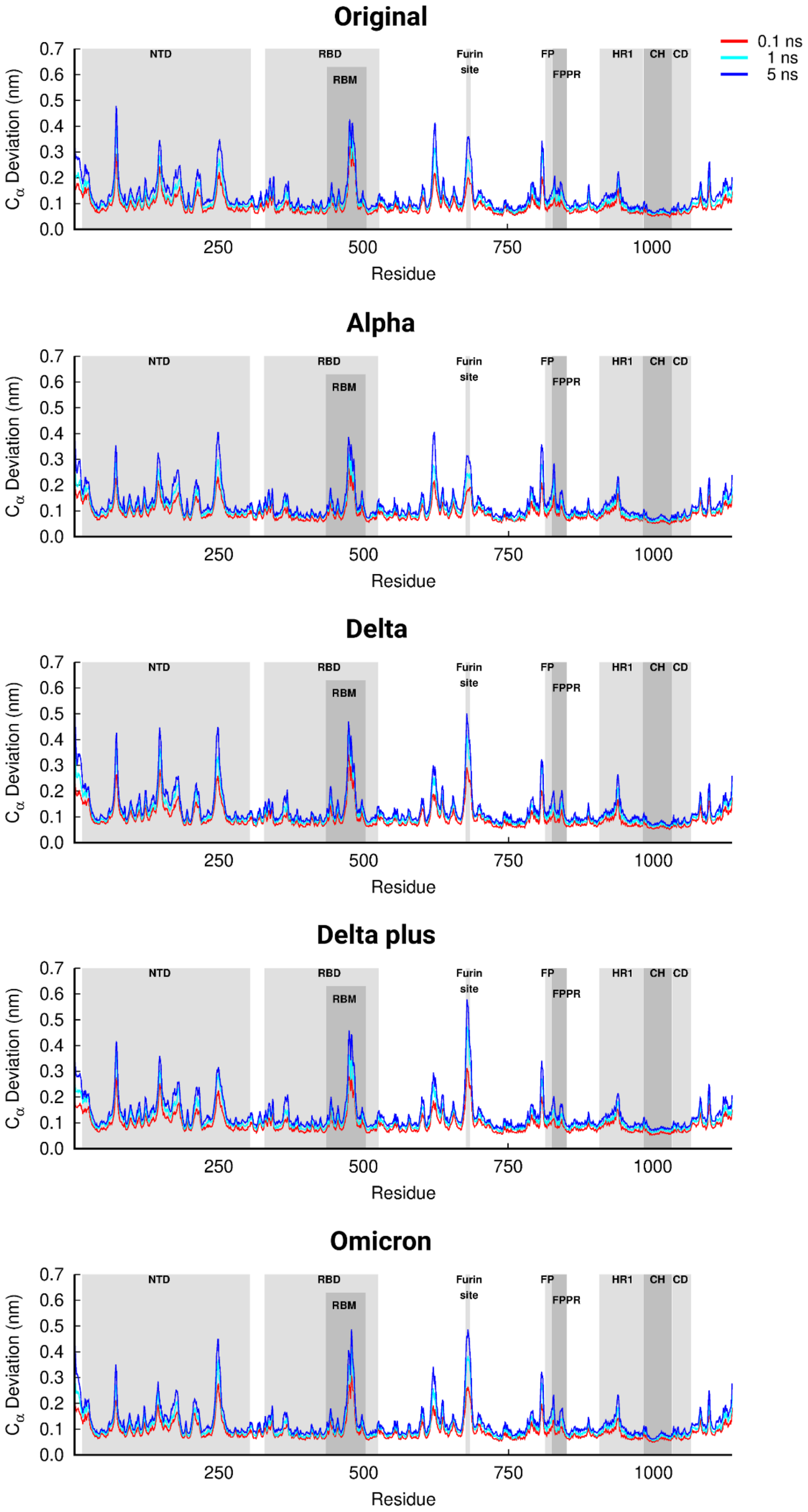
Original
Alpha
Delta
Delta plus
Omicron
0.1 ns
1 ns
5 ns
NTD
RBD
RBM
Furin site
FP
FPPR
HR1
CH
CD
Cα Deviation (nm)
Residue
0.0
0.1
0.2
0.3
0.4
0.5
0.6
0.7
250
500
750
1000

**Figure S3.** Average Cα-positional deviations in the five nanoseconds after LA removal from the FA sites in the original, Alpha, Delta, Delta Plus and Omicron spikes. The average deviations were determined using the Kubo-Onsager approach (*13-16*) for the pairwise comparison between the D-NEMD apo and equilibrium LA-bound simulations. The averages were calculated over the three chains of the trimer and over 90 pairs of simulations. The positions of some important structural motifs are highlighted in grey, namely the N-terminal domain (NTD), receptor-binding domain (RBD), receptor-binding motif (RBM), fusion peptide (FP), fusion-peptide proximal region (FPPR), heptad repeat 1 (HR1), central helix (CH) and connector domain (CD). Please zoom in to the image for detailed visualisation.

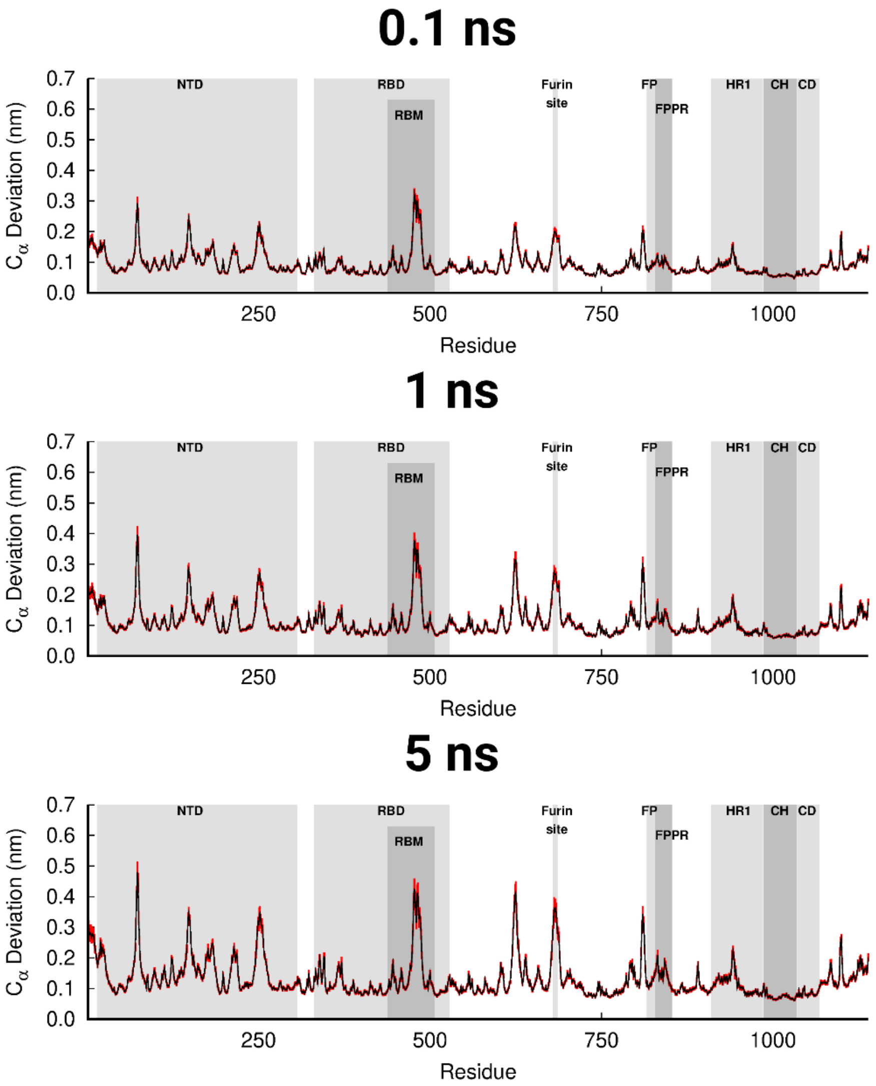


**Figure S4.** Average Cα-positional deviations and corresponding standard errors in the 0.1, 1 and 5 ns after LA removal from the FA sites in the original spike protein from SARS-CoV-2. The average deviations were determined using the Kubo-Onsager approach (*13-16*) for the pairwise comparison between the nonequilibrium apo and equilibrium LA-bound simulations. The averages were calculated over the three chains of the trimer and over 90 pairs of simulations. The vertical red lines represent the standard error of the mean. The positions of some important structural motifs are highlighted in grey, namely the N-terminal domain (NTD), receptor-binding domain (RBD), receptor-binding motif (RBM), fusion peptide (FP), fusion-peptide proximal region (FPPR), heptad repeat 1 (HR1), central helix (CH), connector domain (CD). Please zoom in to the image for detailed visualisation.

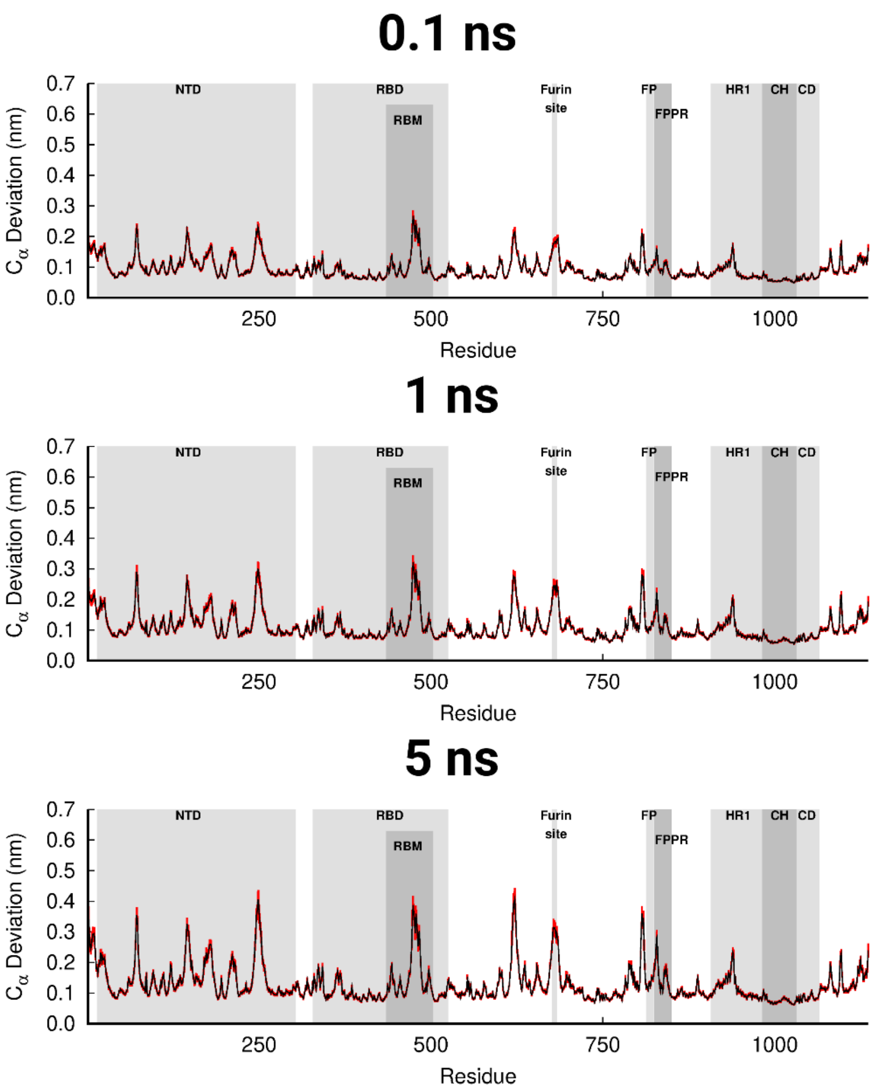


**Figure S5.** Average Cα-positional deviations and corresponding standard errors in the 0.1, 1 and 5 ns after LA removal from the FA sites in the Alpha spike protein from SARS-CoV-2. For more details, see the legend in Figure S4. Please zoom in to the image for detailed visualisation.

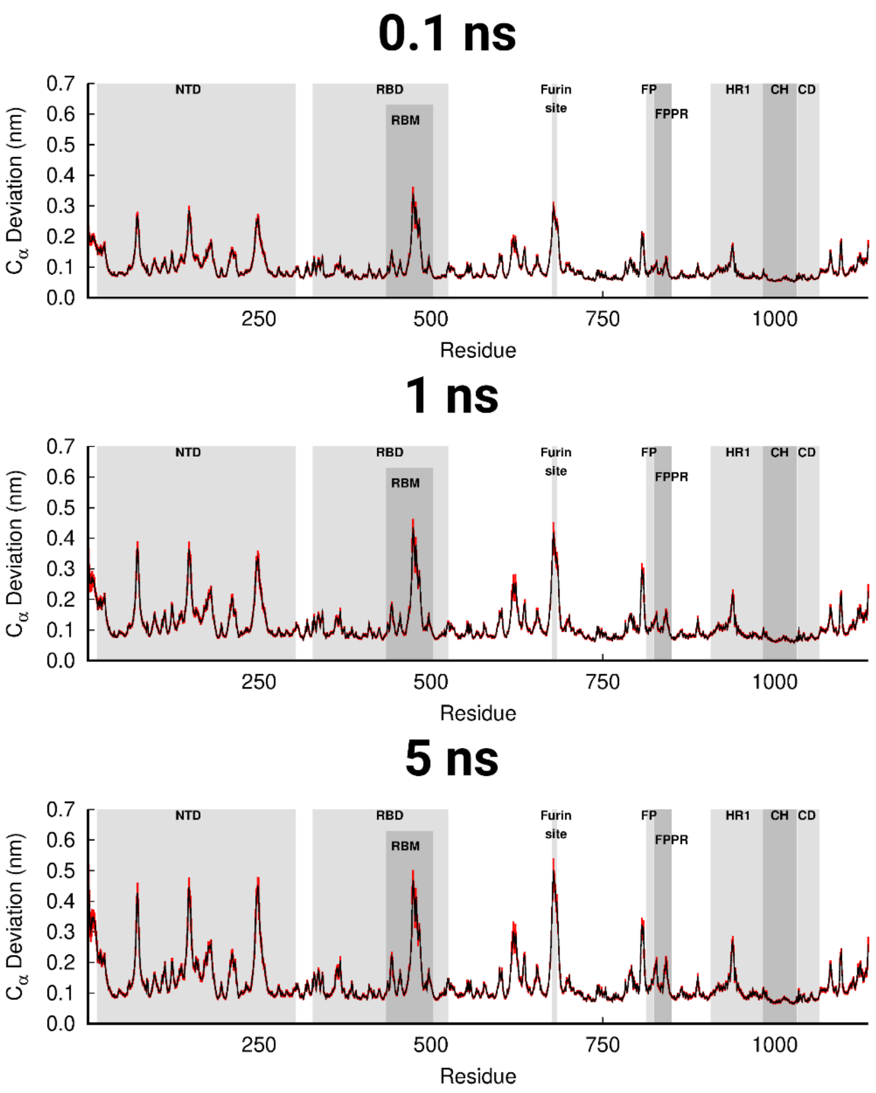


**Figure S6.** Average Cα-positional deviations and corresponding standard errors in the 0.1, 1 and 5 ns after LA removal from the FA sites in the Delta spike protein from SARS-CoV-2. For more details, see the legend in Figure S4. Please zoom in to the image for detailed visualisation.

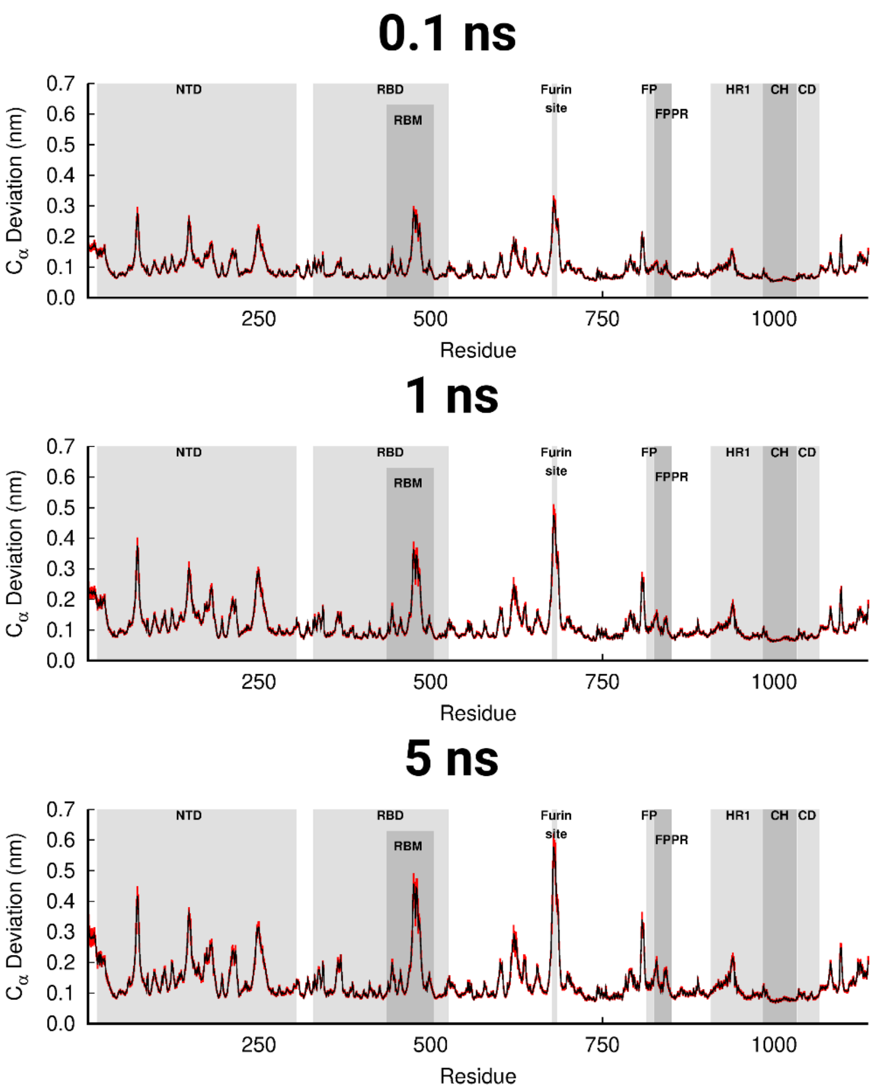


**Figure S7.** Average Cα-positional deviations (and corresponding standard errors) in the 0.1, 1 and 5 ns after LA removal from the FA sites in the Delta Plus spike protein from SARS-CoV-2. For more details, see the legend in Figure S4. Please zoom in to the image for detailed visualisation.

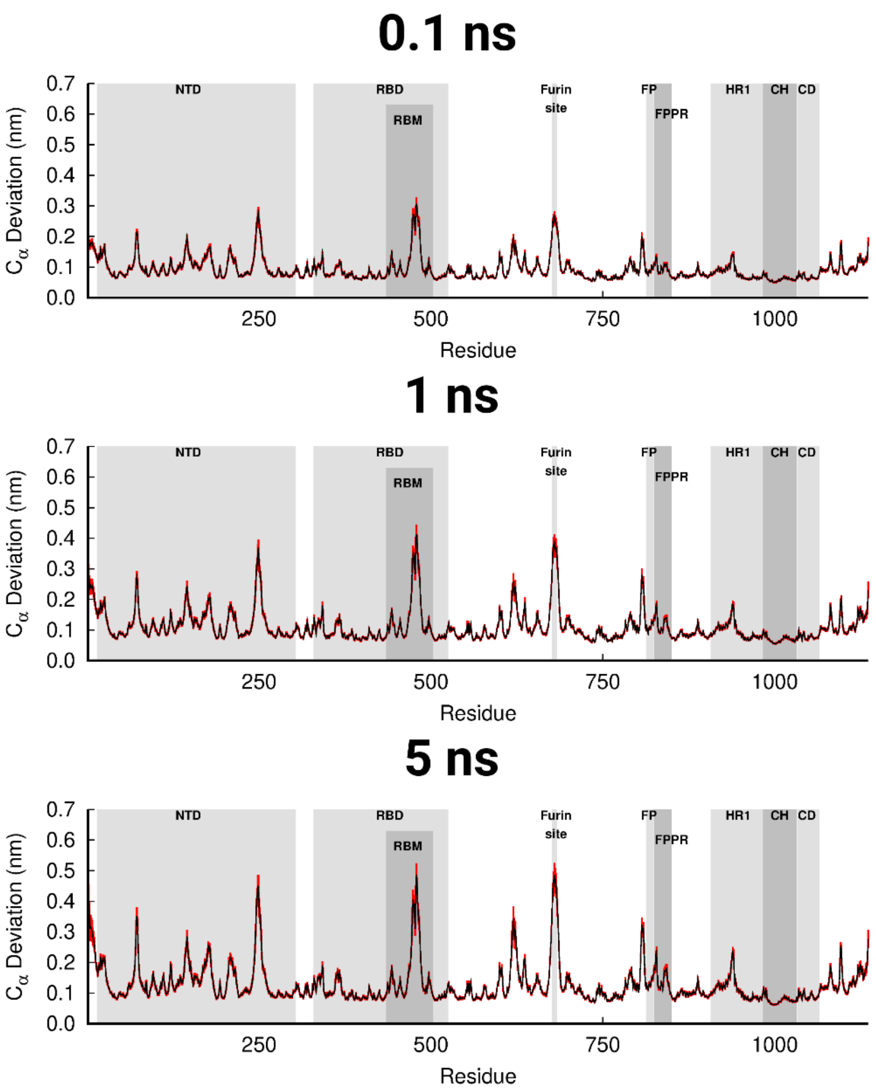


**Figure S8.** Average Cα-positional deviations (and corresponding standard errors) in the 0.1, 1 and 5 ns after LA removal from the FA sites in the Omicron spike protein from SARS-CoV-2. For more details, see the legend in Figure S4. Please zoom in to the image for detailed visualisation.

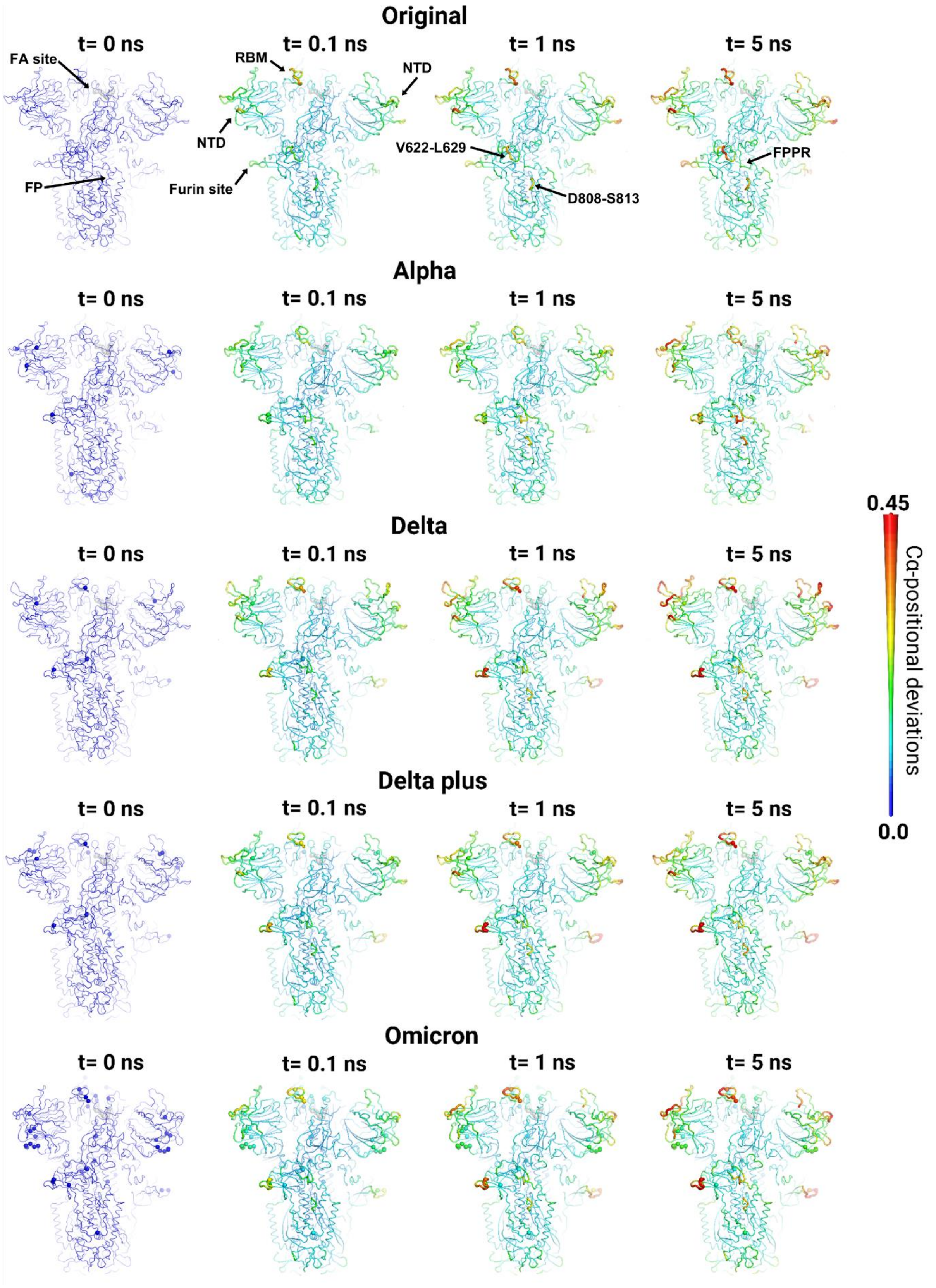


**Figure S9.** Mapping of the average Cα-positional deviations in the five nanoseconds following LA removal in the original, Alpha, Delta, Delta Plus and Omicron spike. The Cα deviations between the D-NEMD apo and equilibrium LA-bound simulations at specific times (0, 0.1, 1 and 5 ns) after LA removal were calculated as a function of the residue number. The final deviation values correspond to the average obtained over the three chains of the trimer and over all 90 pairs of simulations (Figure S3). The Cα average deviations are mapped onto the structure used as the starting point for the LA-bound equilibrium simulations of each variant. Note that in this image, both structure colours and cartoon thickness relates to the average Cα-positional deviation values. The grey spheres highlight the location of the FA binding site. The remaining spheres pinpoint the position of mutations. Please zoom in to the picture for detailed visualisation.

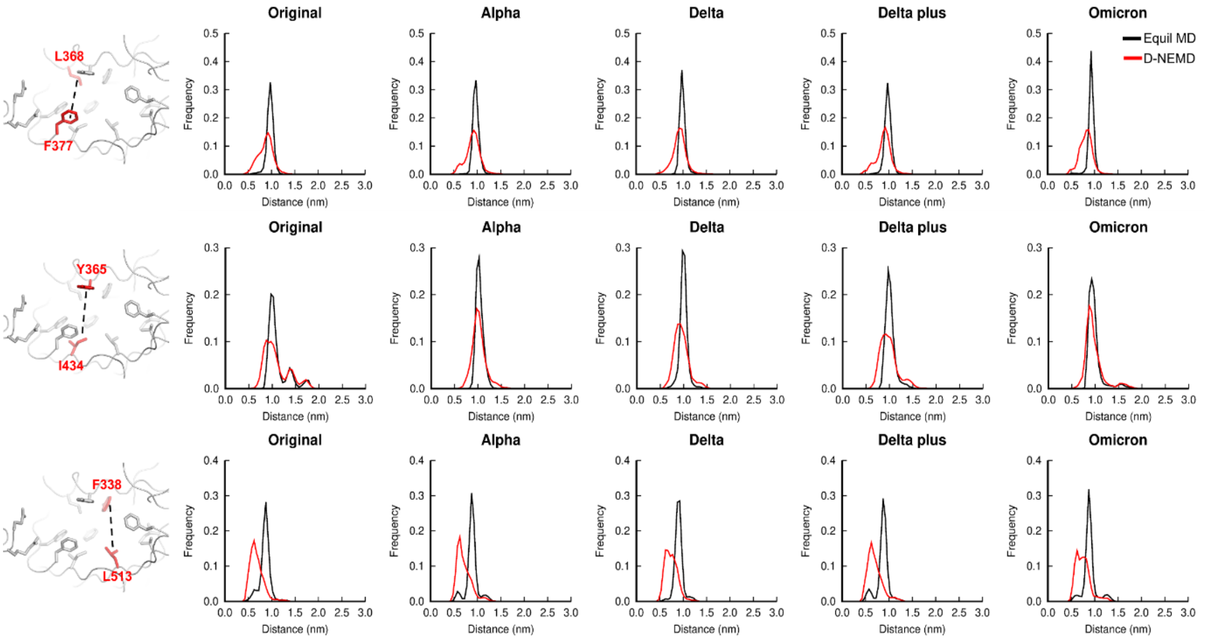


**Figure S10.** Distributions of the distances between L368-F377, Y365-I434 and F338-L513 in the equilibrium FA-bound and D-NEMD apo original, Alpha, Delta, Delta Plus and Omicron variant. Overall distribution of the distance between the centre of mass of the sidechains of L368 and F377, Y365 and I434 and F338 and L513 in the equilibrium (black line) and D-NEMD (red line) simulations. Note that the images of the FA site (shown on the left) represent the same orientation as Figure 1B. Please zoom in to the image for detailed visualisation.

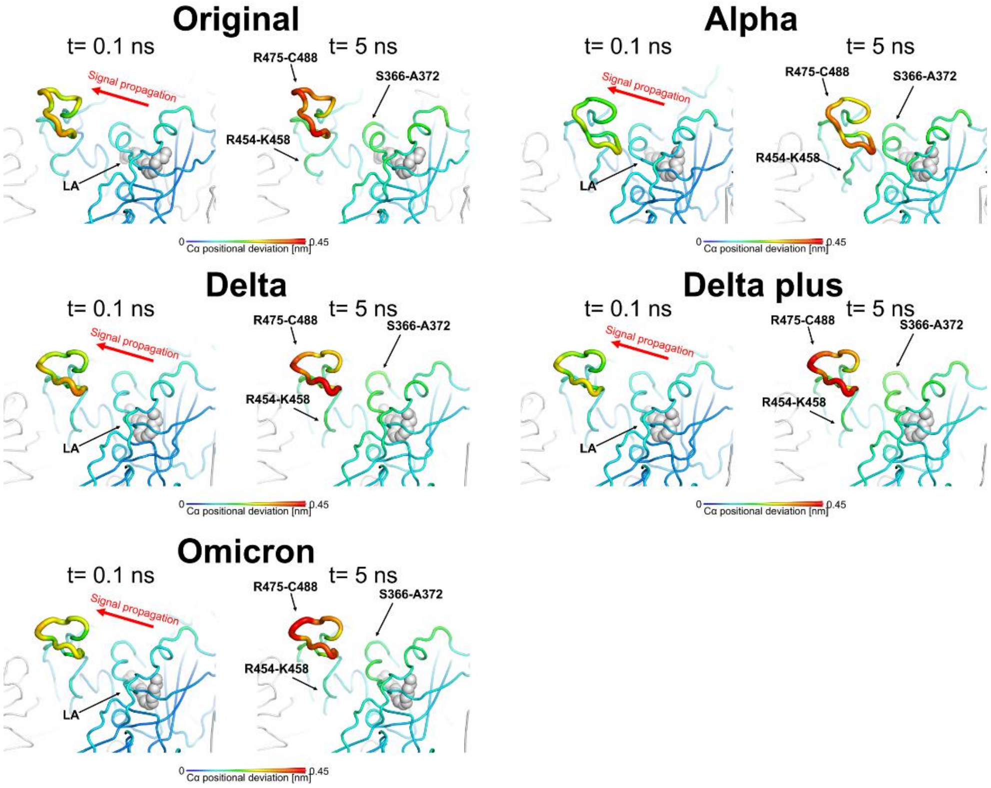


**Figure S11.** Apparent allosteric pathway connecting the FA site to the RBD in the original, Alpha, Delta, Delta plus and Omicron. Average Cα-positional deviation at times 0.1 and 5 ns following LA removal from the FA binding pockets. For more details, see the legend in Figure S9. Please zoom in to the picture for detailed visualisation.

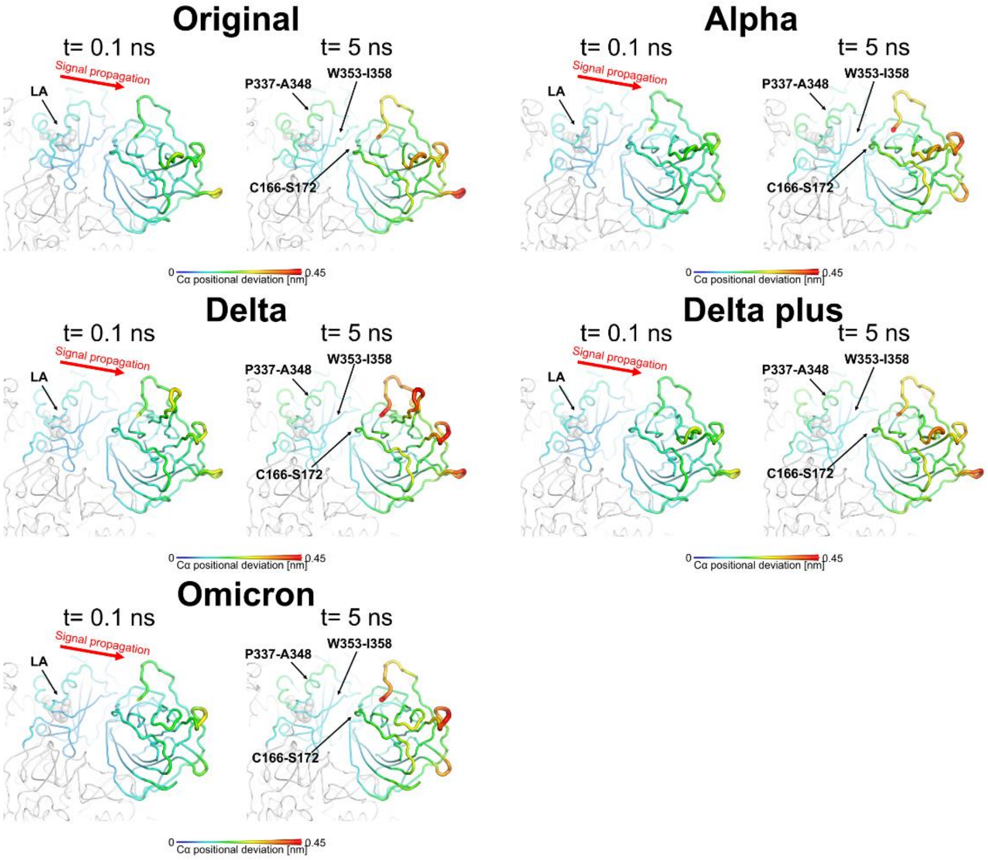


**Figure S12.** Apparent allosteric pathway connecting the FA site to the NTD in the original, Alpha, Delta, Delta plus and Omicron. Average Cα-positional deviation at times 0.1 and 5 ns following LA removal from the FA binding pockets. For more details, see the legend in Figure S9. Please zoom in to the picture for detailed visualisation.

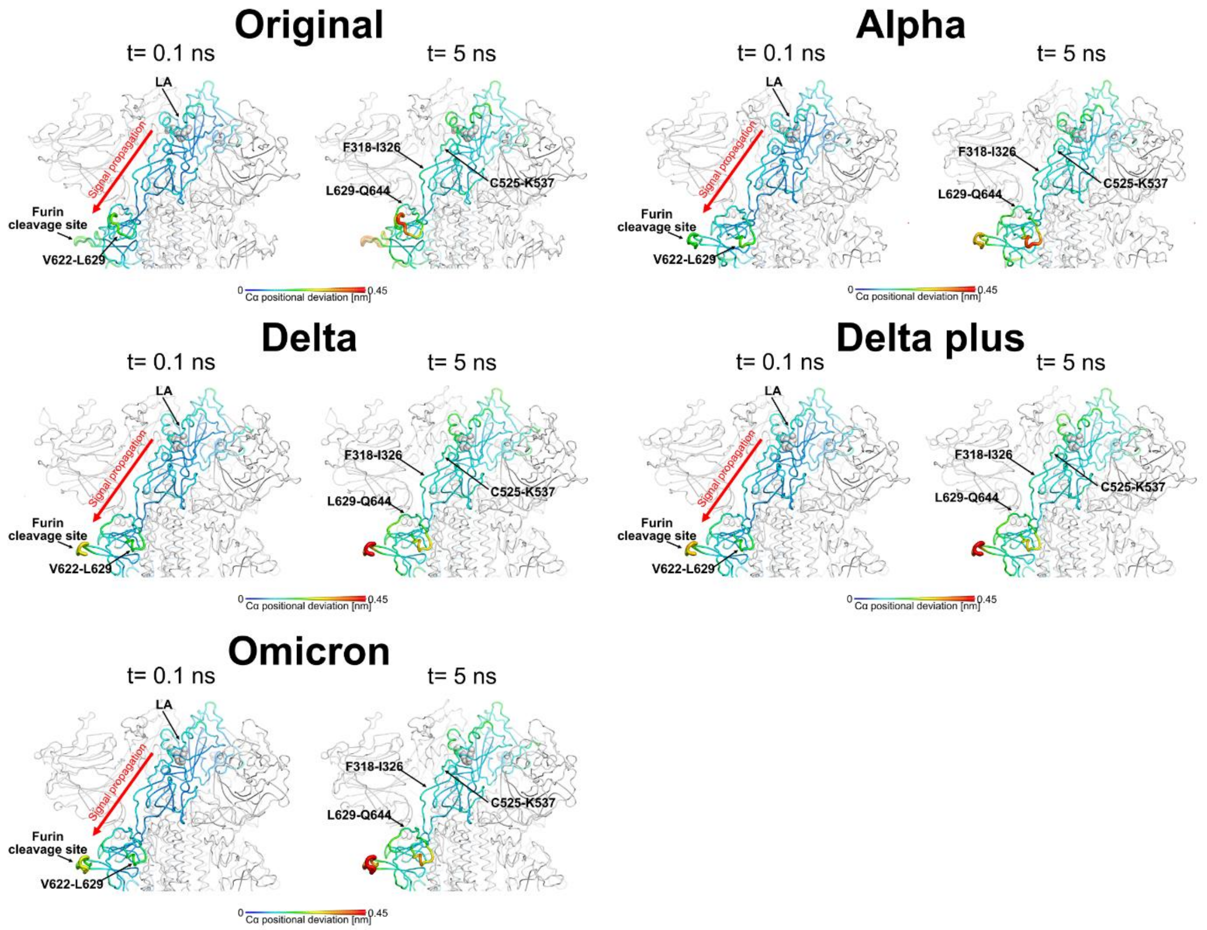


**Figure S13.** Apparent allosteric pathway connecting the FA site to the furin cleavage region in the original, Alpha, Delta, Delta plus and Omicron. Average Cα-positional deviation at times 0.1 and 5 ns following LA removal from the FA binding pockets. For more details, see the legend in Figure S9. Please zoom in to the picture for detailed visualisation.

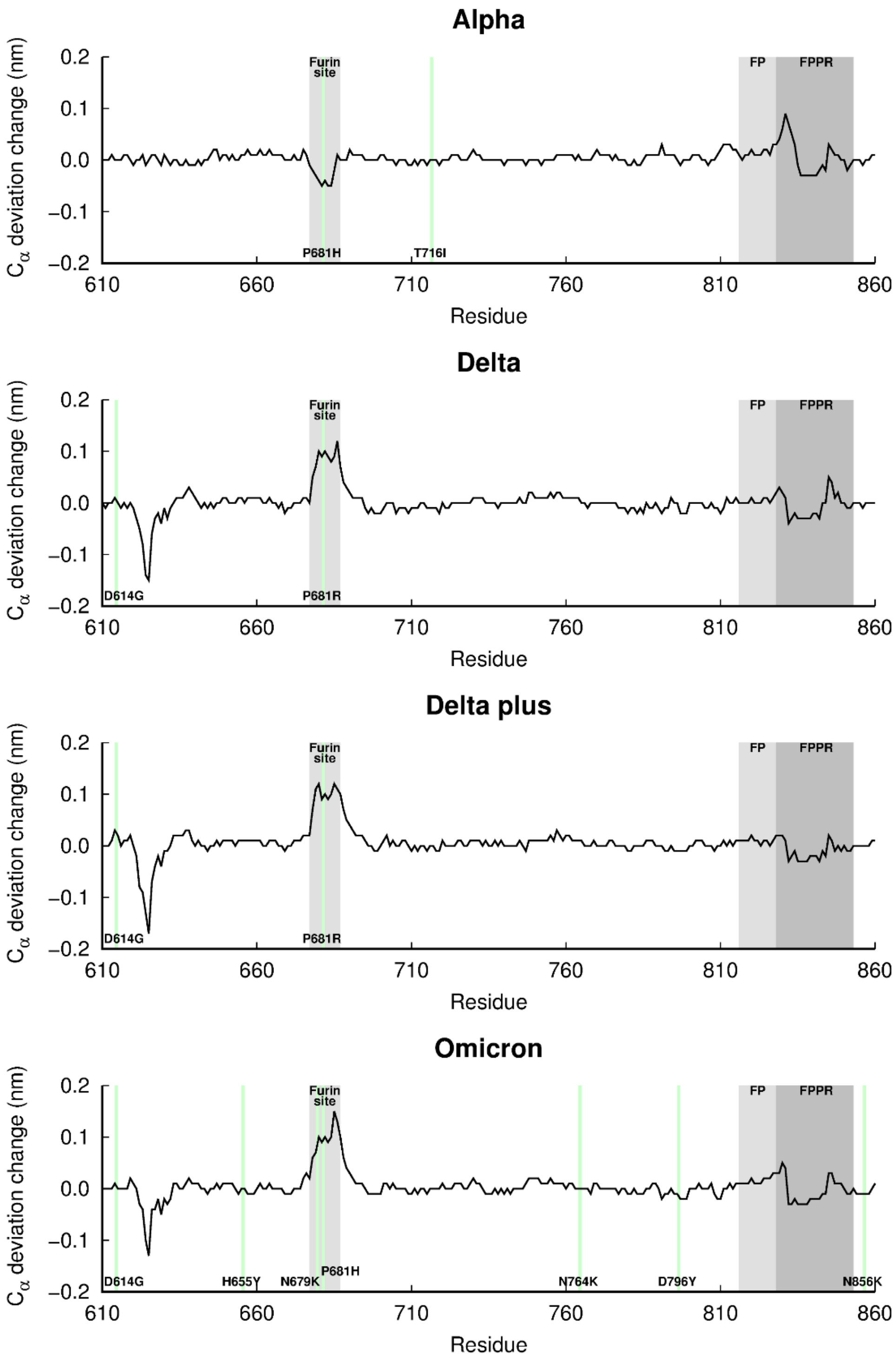


**Figure S14.** Difference in the structural response of the furin cleavage site and FP-surrounding regions between the original and Alpha, Delta, Delta plus and Omicron variants in the t=5 ns after LA removal from the FA sites. The position of the furin cleavage site (located at the S1/S2 interface), fusion peptide (FP) and fusion-peptide proximal region (FPPR) is highlighted in grey. The green vertical lines pinpoint the location of the mutations occurring in each variant.
